# A contractile acto-myosin belt promotes growth anisotropy during the early stages of pectoral fin development in zebrafish

**DOI:** 10.1101/512038

**Authors:** Elena Kardash, Hanh Nguyen, Martin Behrndt, Carl-Philipp Heisenberg, Nadine Peyriéras, Marcos Gonzalez-Gaitan

## Abstract

The zebrafish pectoral fin is an *in vivo* model for vertebrate limb formation, well suited to investigate the integration of molecular and cellular dynamics, the results of which translate into shaping the limb bud. We used the ratio between the lengths of the anterior-posterior (AP) and dorso-ventral (DV) axes as the descriptor of how fin shape changes over time. We showed that fin shape transitions from close to hemi-spherical (ratio 1. 36 ± 0.11) to semi-ellipsoid (ratio 1.64 ± 0.04) between 33 and 46 hours post fertilization (hpf). This shape transition coincided with the formation of a contractile “actin belt” at the distal rim of the fin bud along its AP axis. The actin belt emerged from a central position and expanded on both sides along the distal rim of the fin, thus marking the DV boundary between two rows of ectodermal cells. Formation of the actin belt depended on Rac protein activity, as suggested by FRET measurements using a Rac biosensor. 3D+time imaging of the developing fin in Rac-deficient embryos showed that anisotropic growth of the fin depends on the actin belt. Indeed, actin belt formation was dramatically reduced or even absent in the embryos without proper Rac activity. This correlated with isotropic growth of the fin bud from normal shape at 33 hpf to quasi hemispherical shape with AP/DV ratio ~1 13 hours later, without affecting cell number and overall bud volume. We propose that the formation of a contractile acto-myosin belt is essential to drive the pectoral fin’s early anisotropic growth.

## Results and discussion

### *mPrxl(cFos):EGFP* transgenic line reveals two different cellular populations in the mesodermal portion of the fin tissue

The pectoral fin of the zebrafish serves as an *in vivo* model for vertebrate limb development [1]. The fin field is specified by the expression of the transcription factor Tbx5a in the lateral plate mesoderm (LPM) at 18 hours post fertilisation (hpf) [2]. Most studies on the pectoral fin are focused on the late stages of its development (after 36 hpf or later), when the fin is already morphologically distinct [3, 4]. However, little is known about the mechanisms that control the establishment of fin shape during the early stages of its development. Using the *Et(hand2:EGFP)ch2* transgenic lines, in which all LPM cells are labeled, Mao *et al.* showed that LPM cells converge and subsequently undergo condensation at the fin field between 18 and 23 hpf [5]. The transcription factor Hand2 is expressed initially throughout the presumptive fin or limb field during the onset of bud outgrowth, but at the later stages it is restricted to the posterior portion of the fin mesenchyme ([6–8]. Therefore, only posterior fin cells are labelled in *Et(hand2:EGFP)ch2* transgenic embryos after the onset of fin formation. To monitor all fin-specific LPM cells, including both the anterior and posterior domains during the first 48 hours of development, we used a recently established transgenic line *Tg(mPrx1(cFos):EGFP)*, in which fin mesenchyme is labelled by EGFP expression, driven by the mouse *prx1* enhancer from 26 hpf onwards[9]. Starting from 26 hpf, we observed two broad domains of EGFP signal on both sides of the notochord, corresponding to the area of LPM adjacent to 2^nd^ and 3^rd^ somites (Figure 1A). After 30 hpf, the fin mesodermal core comprised two populations of cells: EGFP-positive and EGFP-negative cells (Figure 1B, sagittal and lateral views). Live imaging showed an influx of cells into the fin field from the posterior region corresponding to the 4^th^ somite area between 32 and 46 hpf (Movies 1 and 2). Somitic mesodermal cells migrate into the fin field from the area corresponding to 4^th^-7^th^ somites, and some contribute to the apical ectodermal ridge (AER) [4]. These invading somitic cells do not express EGFP. Cell tracking showed that these cells indeed formed EGFP-negative domains within the fin tissue (Supplemental Figure 1, Movie3). Our data thus uncover at least two different cellular populations in the fin mesenchyme: the EGFP-positive cells that report the activity of the *mPrx1* enhancer, which labels locally proliferating lateral plate mesoderm cells, and the EGFP-negative invading somitic cells.

**Figure 1.**
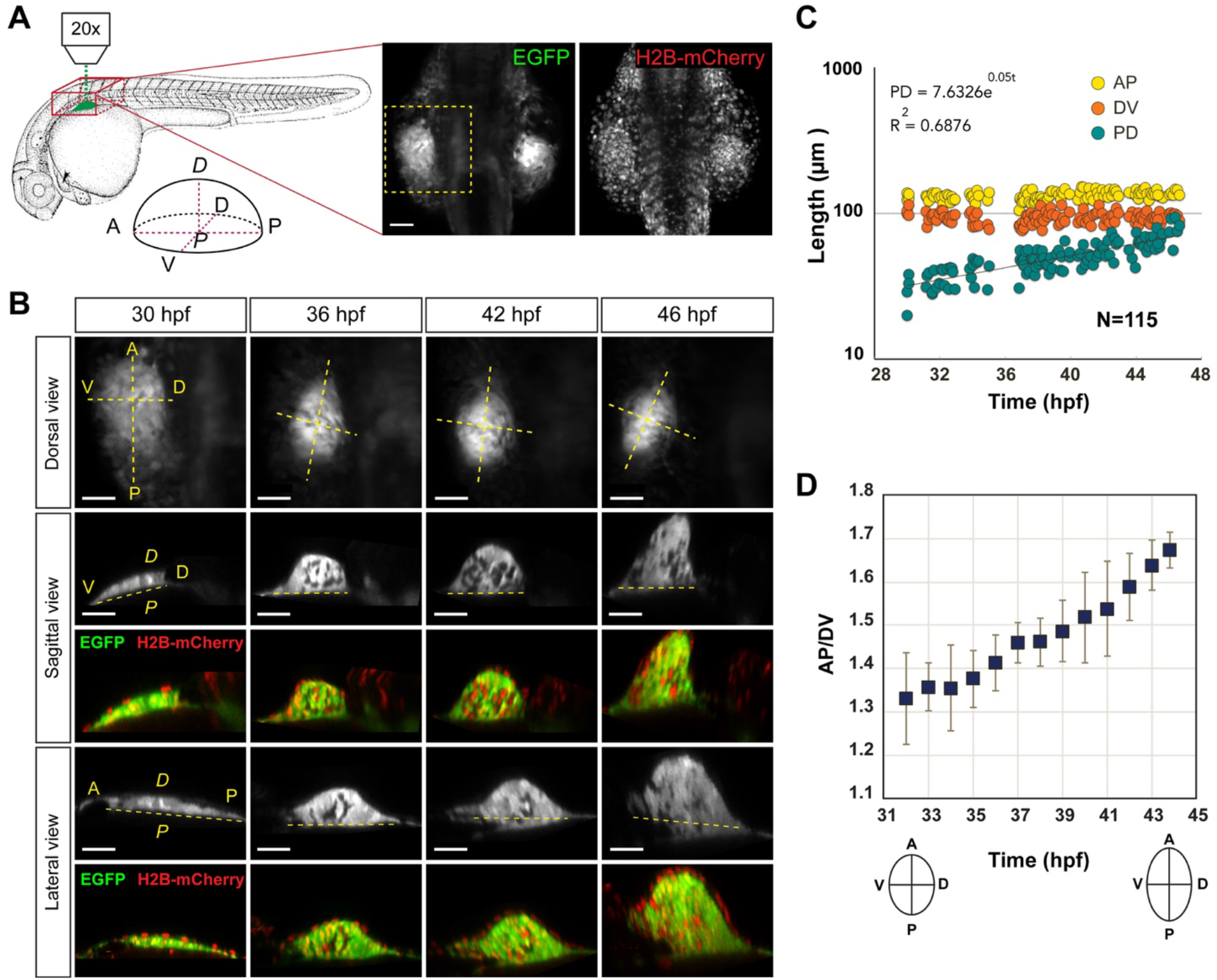
Characterizing the early stages of pectoral fin growth. **A.** The scheme shows our imaging setup to monitor fin growth. The drawing depicts a zebrafish embryo at 31 hpf with the emerging fin labeled in green. Dorsal view shows z projection of a double transgenic line *Tg*(*Ef1a:H2B-mCherry*); *Tg*(*mPrx1 (cFos):EGFP*) at 30 hpf. A and P refer to the anterior and posterior of the embryo. The yellow insert indicates the location of the fin. Scale bar: 50μm. **B.** Representative images of pectoral fins at different developmental stages ranging from 30 to 46 hpf are shown. On the dorsal view, the z projection is shown. A – anterior, P – posterior, D – dorsal, V – ventral, *P* – proximal, *D* – distal. Scale bar: 40μm. **C.** The graph shows the dimensions of each axis of the fin presented as a function of time on a logarithmic scale. N =115. **D.** The graph shows the dynamics in AP/DV ratio in *Tg*(*mPrx1(cFos):EGFP*) embryos between 32 and 44 hpf. N = 6. Data are represented as mean ± standard deviation. See also Supplemental Figure 2A.

### Emerging dimensions and geometry of the pectoral fin bud during early stages of development

Applying the same principles used for the limb bud in birds and mammals [10, 11], we assigned the three principal axes of the fin: anterior-posterior (AP), dorso-ventral (DV), and proximal-distal (*PD*), (Figure 1A). We analysed how fin shape evolves by measuring the dimensions along its principal axes over time. AP and DV axes define the base of the fin, an important structure that connects the fin to the embryo’s body. In the early stages, the EGFP-positive domain appeared more scattered as compared to the later stages, when we saw compaction of the fin field (Figure 1B dorsal view, Supplemental movies 4 and 5). At 36 hpf, the fin bud closely resembled a semi-ellipsoid, based on the observed symmetry along the AP, DV and *PD* axes (Figure 1B). The semi-ellipsoid symmetry in the fin persisted until 46 hpf (Figure 1B, Supplemental movie 5). As expected, major growth occurred along the *PD* axis of the fin (34.7 ± 6 μm at 30 hpf and 72.2 ± 12 μm by 46 hpf) with slight fluctuations in length observed along AP and DV axes (Figure 1C). A closer look showed an interesting relation between the AP and DV axes within the time frame between 32 and 44 hpf. Specifically, during that time, the DV axis shortened from 85 ± 5 μm to 79 ± 5 μm while the AP axis elongated from 116 ± 10 μm to 135 ± 12 μm. We measured changes in AP/DV ratio between 32 and 44 hpf, which showed an increase in the AP/DV ratio from 1.36 ± 0.11 to 1.64 ± 0.04 (Figure 1D, Supplemental Figure 2A, n=6).

### An actin belt is formed in the ectodermal portion of the fin along the anterior-posterior axis, marking its dorso-ventral boundary

Coordinated elongation and shortening of the AP and DV axes, respectively, between 35 and 46 hpf resulted in anisotropic growth of fin tissue. What could cause such shape transition? One possibility is the mechanical stress generated in the growing fin tissue, as shown for mouse limb bud growth, where anisotropic stress in limb ectoderm was observed [12]. To study the biomechanical components of pectoral fin tissue, we first monitored the dynamics of filamentous actin in the fin by visualising Lifeact-RFP under the control of the *actb1* promoter. We discovered the presence of a dense actin structure in the distal rim of the fin along its AP axis (Figure 2A). We named this structure the “actin belt”. Phalloidin staining of endogenous F-actin in at 46 hpf confirmed the presence of the same structure in wild-type embryos (Supplemental Figure 2B). Confocal imaging at high magnification showed that actin accumulated in patches in a polarised fashion, thus creating a dense filamentous network restricted to the two rows of ectodermal cells that mark the DV boundary of the fin (Figure 2A, B and supplemental movie 6). According to its positioning in the ectoderm at the dorso-ventral fin boundary, actin belt is localised to the apical ectodermal ridge (AER), the major signalling centre that controls fin growth. Accumulation of the actin belt was detected at around 35 hpf at the distal portion of the fin. This was followed by actin propagation along the entire AP axis of the fin, creating an arc in the ectodermal layer by 40 hpf (Figure 2C, Supplemental movie 7). Accumulation of this actin belt was transient, disassembling gradually after 46 hpf. Additional ectodermal layers formed during the transition from apical ectodermal ridge to apical ectodermal fold (Figure 2B, lateral view) [3]. The ellipsoid symmetry of the fin was lost at around 48 hpf, as its distal portion bent towards the embryonic body (Supplemental movie 8).

**Figure 2.**
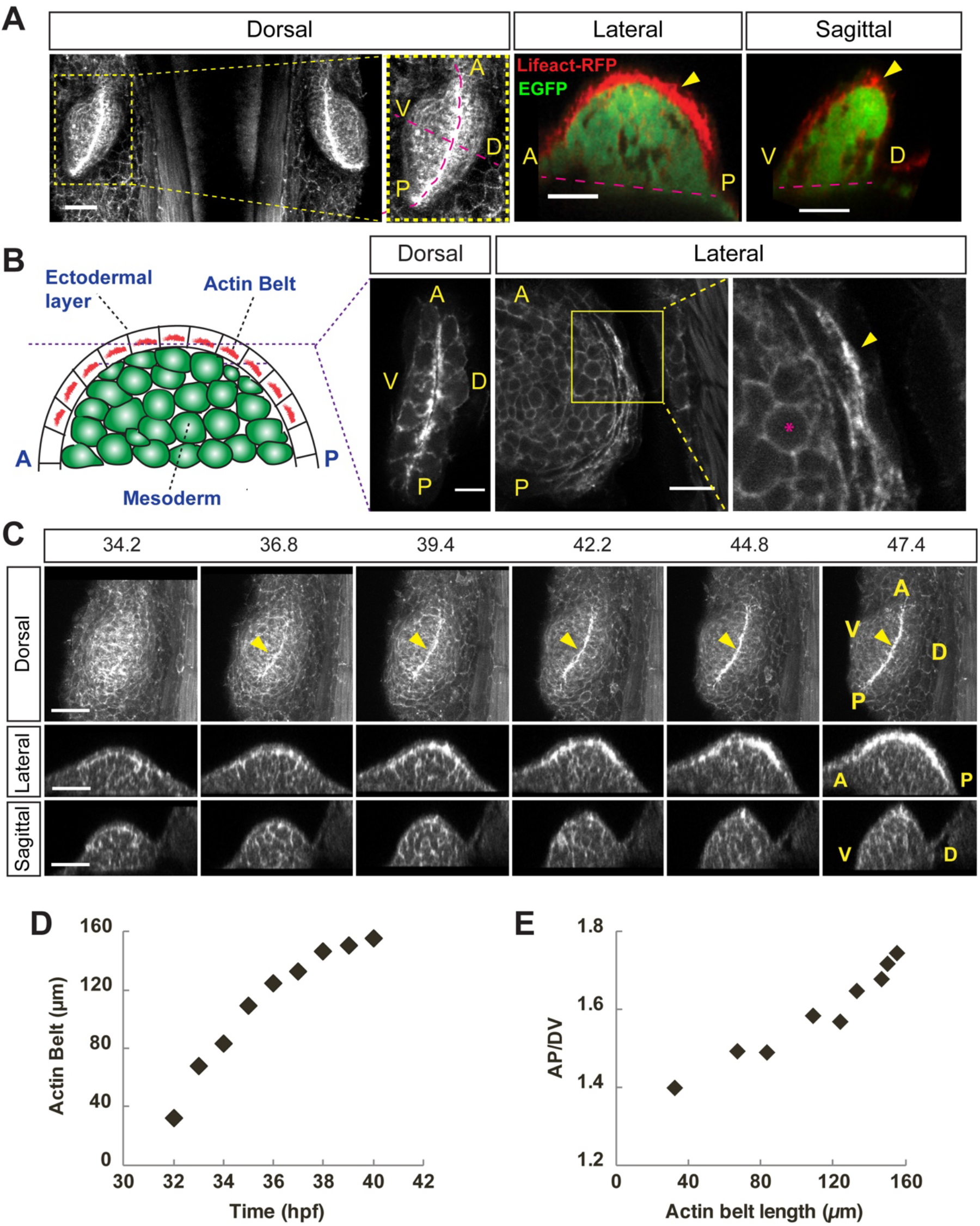
Actin cytoskeleton dynamics during fin formation. **A.** Z projection along the PD shows a dorsal view of the fins in a transgenic line expressing Lifeact-RFP. The lateral and sagittal views show resliced images along the AP and DV axes respectively. A and P refer to the anterior and posterior of the embryo. Scale bar: 40μm. **B.** A scheme shows the cross-section of the pectoral fin along its AP axis at the dorso-ventral boundary. The dorsal view shows a 2μm thick confocal section of the distal part of a fin in a transgenic line expressing Lifeact-RFP. **F.** The lateral view shows a cross-section of the fin along its AP axis at the DV boundary Scale bar: 20μm. Scale bar: 20μm. Stage: 46 hpf. **C.** Progressive actin accumulation at the DV boundary along the AP axis. Individual snapshots from a time-lapse movie show dorsal view of the fin in a transgenic line expressing Lifeact-RFP. Z projections along the proximal-distal axis shows the dorsal view, the optical crosssections along AP and DV axes show the lateral and the sagittal views respectively. The yellow arrow points at the actin belt. A – anterior, P – posterior, D – dorsal, V – ventral. Scale bar: 40μm. **D.** The graph shows the correlation between the actin belt length and the developmental stage for one embryo. **E.** The graph shows the correlation between the AP/DV ratio and the actin belt length for one embryo. The same embryo as in D.

### The actin belt is essential for anisotropic growth of the pectoral fin

Remarkably, the formation and propagation of the actin belt coincided with the anisotropic shape transition seen in the growing fin between 35 and 46 hpf. This suggested a role of the actin belt in establishing fin shape (Figure 2D and E). To understand the functional significance of the actin belt, we perturbed its formation by manipulating the level of Rac GTPases, because Rac proteins regulate actin polymerisation [13]. We detected elevated Rac activity in the region of the actin belt by FRET ratiometric imaging using a RacFRET biosensor [14] (Supplemental Figure 2C). To address the role of the actin belt in shaping of the pectoral fin more directly, we interfered with the function of endogenous Rac1 and Rac2 proteins, the most likely candidates for promoting actin polymerisation. Rac1 is ubiquitously expressed and plays a role in fusion of fast muscle precursor cells [15], while Rac2 is expressed in the pectoral fin and in leukocytes, where it controls their motility [16–18].

Reducing the levels of either the Rac 1 or Rac2 protein with the appropriate morpholinos disrupted formation of the actin belt and affected both fin growth and shape (Figure 3A, Supplemental Figures 2D). We rescued the *rac1* morpholino-induced phenotype by overexpression of a Rac1-EGFP fusion, thus confirming the role for the Rac1 GTPase in establishing the actin belt (Supplemental Figure 2E). Use of the *rac1* morpholino caused necrotic tissue defects in many embryos. We therefore conducted subsequent experiments with the *rac2* morpholino, which gave rise to fewer side effects owing to its restricted expression in pectoral fins and neutrophils [17, 18]. We observed two main categories in the *rac2* morphants: i) a severe phenotype: the actin belt was missing entirely and the fins were of smaller size (Supplemental Figure 3A), and ii) a partial phenotype: a partially formed actin belt with fins of similar size as control fins (Figure 3A and Supplemental Figure 3 A’, A’’ and B). In all fins where the actin belt was missing, punctured or reduced, the AP/DV ratio was reduced as well (Figure 3B, C). Rac2-deficient fins with a partial actin belt phenotype had the same number of cells as compared to control fins at 46 hpf. This showed that in embryos with this partial phenotype cell proliferation in the fin mesenchyme was not affected (Supplemental Figure 3B). Similarly, the migration of the somitic mesodermal cells into the fin area appeared unaffected in fins with impaired actin belt in the embryos with a partial phenotype. This conclusion was supported by the presence in *rac2* morphant fins of EGFP-negative cells on the *mPrx1(cFos):EGFP* transgenic background at 46 hpf and their migration into the fin area (Figure 3A and supplemental Figure 3A’ and A’’, Supplemental movie 9).

**Figure 3.**
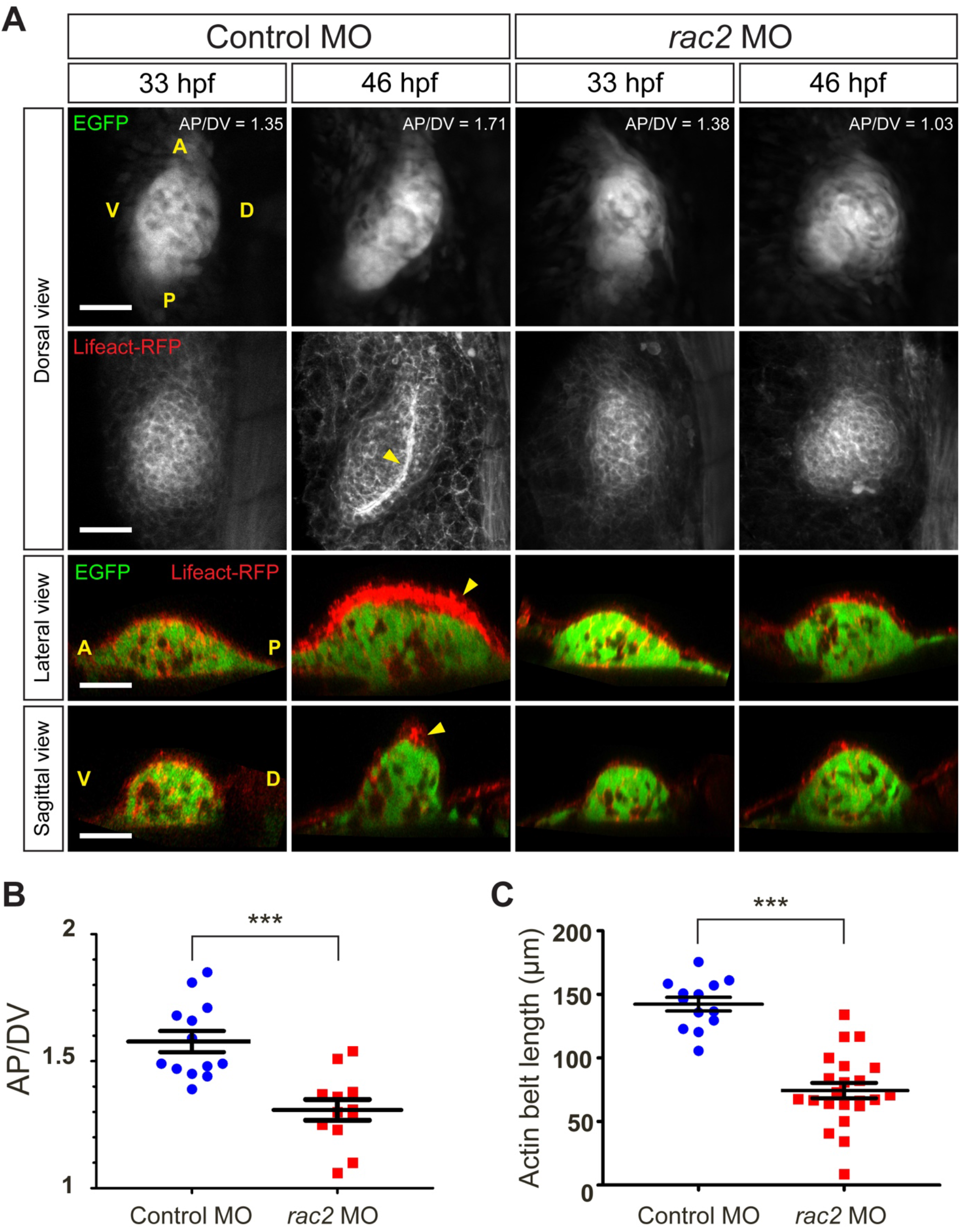
The role of Rac2 in actin belt formation and fin shape establishment. **A.** Pectoral fins in double transgenic embryos *Tg*(*mPrx1(cFos):EGFP*); *Tg*(*actb1:Lifeact-RFP*) were injected either with the control or *rac2* morpholinos and the images of the fin were taken at 33 and 46 hpf. The yellow arrow points at the actin belt. Scale bar: 40μm. **B.** The graph compares the AP/DV ratios in the fins of the control and *rac2* morpholino treated embryos at 46 hpf. N=13 for the control and n=12 for the Rac2MO treated embryos. Data are represented as mean ± SEM. *p* = 0.0006. See also Figure S3. **C.** The graph compares actin belt length at 46 hpf in fins of the control and *rac2* morpholino treated embryos. N= 13 for the control and n=22 for the *Rac2MO*. Error bars represent SEM. *p* < 0.0006. See also Figure S3.

Both *rac1* and *rac2* morpholinos had similar effects on formation of the actin belt and in establishing fin shape, suggesting a functional redundancy of Rac1 and Rac2. Our findings agree with an earlier suggestion that Rac1 and Rac2 can have partially overlapping functions *in vivo* [18]. Reducing the level of either one of these proteins was sufficient to cause the loss of the actin belt in pectoral fins. While Rac1 and Rac2 proteins might both contribute to actin polymerisation during formation of the belt, a minimal level of Rac activity is likely required for proper assembly of the actin belt.

### Dynamics of cellular behavior during fin growth and actin belt contractility

Our experiments establish a critical role for the actin belt in the control of anisotropic growth of the pectoral fin pectoral. How could the presence of the actin belt contribute to an anisotropic shape transition? We consider several options: i) the actin belt may generate tension within the fin and favor cell division and cellular rearrangements along the AP axis, thereby causing preferential elongation of the fin along the AP axis; ii) presence of the actin belt may create a physical boundary and cause differential tension along the AP and DV axes. To understand the functional role of the actin belt, we imaged the distribution of myosin, an important actin binding protein that an generate stresses in living tissue. Myosin was indeed enriched at the exact location of the actin belt (Supplemental Figure 4A). The presence of myosin suggested the contractile nature of the actin belt. Indeed, laser ablation of the filamentous actin in the region of the actin belt relaxed this contractility and thus showed strong tension in the actin belt, consistent with stress along the AP axis (Supplemental Figure 4B, Supplemental movie 10).

Fin growth occurs through a combination of cell proliferation in the lateral plate mesoderm, which contributes to fin mesenchyme, and cell migration from the areas of the 4^th^ and 7^th^ somites, which contribute to both fin mesenchyme and to the apical ectodermal ridge, the AER [4, 19]. Oriented cell motility and divisions contribute to shaping the limb bud in the mouse and the pectoral fin in zebrafish [5, 19, 20]. Nonetheless, how cell dynamic behavior contributes to the observed increase in AP/DV ratio remains to be explored (Figure 1D). We therefore followed individual nuclei in the growing fin between 35 to 46 hpf in a transgenic line that expresses a fluorescent histone, H2B-mCherry, under the control of *Xla.Eef1* promoter. We found that most of the cell divisions during that time were aligned along the plane defined by the AP and DV axes of the fin, with a slight bias towards the PD axis for a few cells. (Figure 4A). The orientations of mitotic divisions appeared random, with a slight bias towards the AP axis of the fin (Figure 4A). These observations suggest that at the early stage of fin growth, mitotic orientations do not obviously contribute to the establishment of its shape, as opposed to what has been shown for mouse and chick limbs [21]. While dividing cells provide the bulk material to build the fin, it is not yet clear how shape of the fin is controlled.

**Figure 4.**
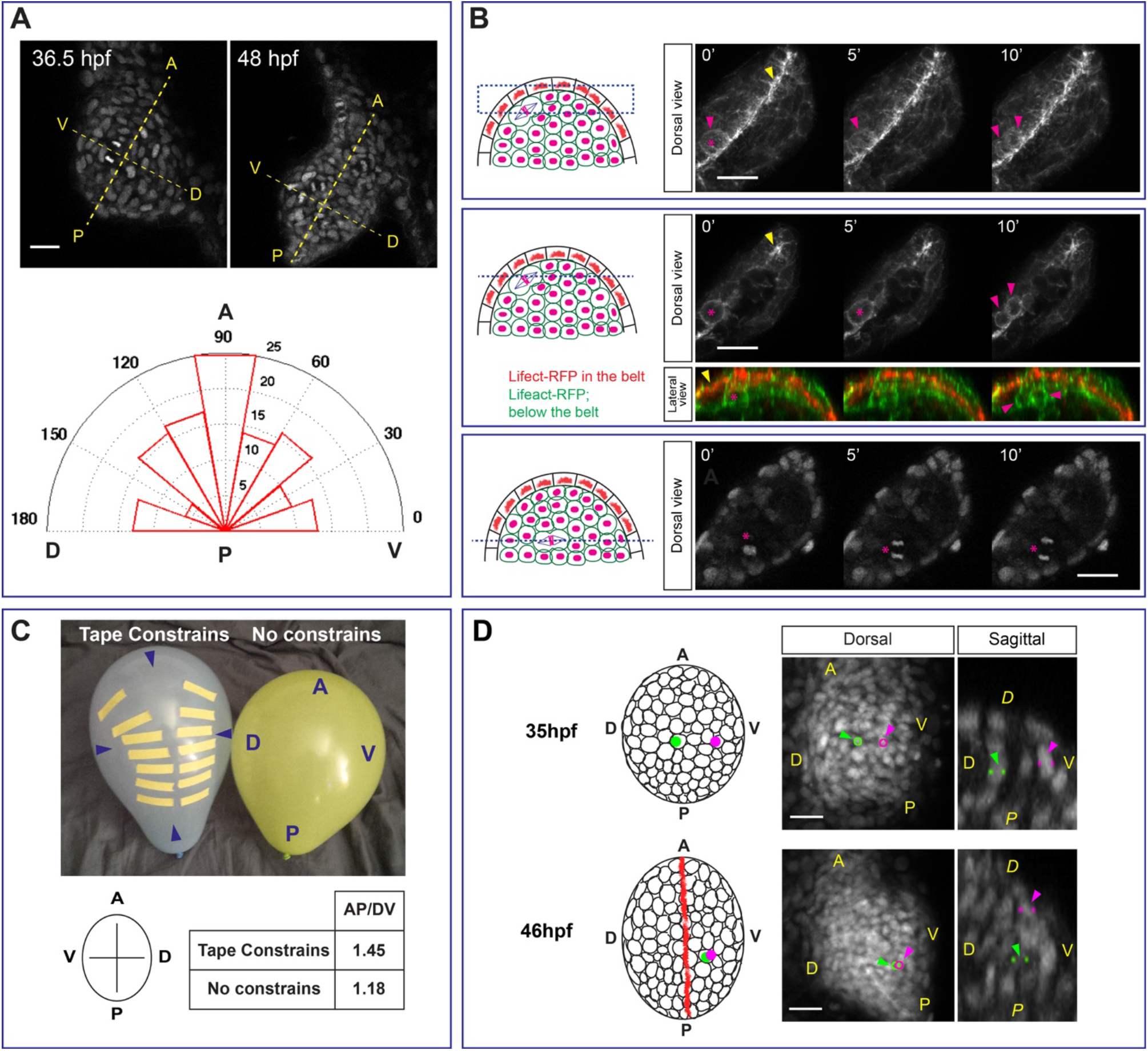
Cellular dynamics during anisotropic fin growth. **A.** Confocal sections show the planes of the fin defined by the AP and DV axes in a *Tg*(*X1a.Eef1α:H2B-mCherry*) embryo. Mitotic orientations angles were measured as projections onto the plane defined by AP and DV axes and relative to the DV axis. Scale bar: 20μm. A polar plot summarizes angles for mitotic orientations between 34 and 49 hpf. Bin 18, n=130. **B.** The top panel shows a z-projection of the dorsal portion of the fin in embryos expressing Lifeact-EGFP. The middle panel shows a single confocal section through the dividing cell. In the lateral view, the actin in the ectodermal layer is labelled with red and the actin in the mesodermal layer is labelled in green. The asterisk and the pink arrow indicate the mesodermal cell undergoing division, the yellow asterisk points at the actin belt. The bottom panel shows a confocal section from the deeper layer of fin mesoderm, the asterisk is next to the cell undergoing division. Double transgenic embryos *Tg*(*beta-actin:Lifeact-GFP*); *Tg*(*X1a.Eef1α:H2B-mCherry*) are at 46 hpf. Scale bars: 40μm. **C.** Two identical balloons were inflated either with or without adhesive tape constrains. The grey balloon on the left was subjected to adhesive tape constrains at the early stages of inflation and mimics the wild-type fin, the yellow balloon on the right represents the Rac2-/- fin, that is inflated without constrains. The blue arrows on the grey balloon mark the corresponding AP and DV axes. **D.** The scheme shows dorsal views of the fin at 35 and 46 hpf. The corresponding fin images from a 3D+time confocal sequence of the *Tg*(*X1a.Eef1α:H2B-mCherry*) embryo are shown on the right. Two cells marked in green and magenta were picked randomly at 35 hpf and followed until 46 hpf.

In *Drosophila* epithelium cell divisions align with tissue tension [22]. Similarly, cells in the proximity of the actin belt, close to the tension field, might orient their mitotic spindles along the AP axis and thus cause preferential tissue extension along the AP axis. We noticed that mesodermal cells near the actin belt divide along the AP axis of the fin, while cells found in the deeper portion of the fin divide in random directions (Figure 4B).

### Differential tension along the DV and AP axes of the fin

In the mouse limb bud, a dorso-ventrally-biased stress pattern guides ectodermal remodelling during the formation of the apical ectodermal ridge [12]. In the pectoral fin, the actin belt might generate a physical boundary that separates the dorsal and ventral halves of the fin. This boundary could impose a physical barrier that causes differential tension along AP and DV axes in the fin ectoderm, thus favoring fin elongation along the AP axis. If tension along the DV axes is higher than that along the AP axis, the fin would be forced to extend along the AP axis. We tested this theory of fin growth in a toy model of the fin using balloons. In a balloon, the elastic rubber represents the ectodermal cell layer (as mentioned above, a single ectodermal layer is present in the fin at these stages), while the expanding air corresponds to the expanding mesodermal tissue, which constitutes the inner portion of the fin (Figure 4C). To mimic the physical boundary generated by the actin belt, we placed non-expandable adhesive tape along the AP axis of the balloon and continued to inflate it. While the control balloon without tape expanded isotropically, the balloon with tape expands more along its AP axis than along the DV axis (Figure 4C).

Finally, anisotropic growth of the fin might be controlled by differential adhesion and surface tension between different cells in the fin mesenchyme, thus allowing cells to sort themselves out [23]. Previous work and our 3D+time imaging data showed that cells in the fin mesenchyme have different origins, such as lateral plate mesoderm and somitic mesodermal cells. The latter invade the fin field from the area neighbouring the 4^th^ somite, which likely influences their physical properties. By tracking individual cells, we observed cellular rearrangements within the fin field between 35 hpf, which is the onset of actin belt formation, and 46 hpf, a time at which formation of the actin belt is complete (Figure 4D). The rearrangements observed for the chosen cells are consistent with a pattern that could yield shortening of the DV axis of the fin. To confirm this theory, it would be necessary to address cellular dynamics in full within the entire fin field, an undertaking that would require a massive computational and analytical effort beyond the scope of the present study.

Our results demonstrate the formation of a contractile actin belt localised to the two rows of cells in the ectodermal layer that forms the dorso-ventral boundary of the pectoral fin. We propose a critical role for this actin belt in regulating anisotropic growth of the pectoral fin. Without a properly formed actin belt, the shape of the pectoral fin was compromised, while cell proliferation and migration was mostly unaffected. In addition, our data showed an unexpected and useful property of the *mPrx1(cFos):EGFP* transgenic line, which enabled the study of the different cell populations that contribute to pectoral fin tissue. More work would be required to more precisely determine the exact mechanism by which the actin belt regulates anisotropic fin growth. Next steps in a strategic experimental approach must combine imaging at high spatio-temporal resolution with quantitative image processing. Biophysical methods such as laser ablation of cortical actin in ectodermal fin tissue to probe its mechanical properties will be an obvious next step to explore the function of the actin belt in three-dimensional shaping of the zebrafish pectoral fin.

## Experimental Procedures

### Fish husbandry and transgenic lines

Adult zebrafish (*Danio rerio*) were maintained at 28°C according to standard procedures as described in (ref: Kimmel 1995). Embryos were kept at 0.3% Danieau’s medium at 28°C.

The following transgenic zebrafish lines were used in this study: *Tg*(*mPrx1(cFos):EGFP*) line to visualise fin mesenchyme was kindly provided by Carolina Minguillon [9]; *Tg*(*actb1-lifeact-GFP*), *Tg*(*actb1:lifeact-RFP*) and *Tg*(*β-actin::myl12.1-mCherry*) lines to visualise filamentous actin and myosin were a kind gift of Carl-Philipp Heisenberg [24] for filamentous actin and [25] for myosin labelling respectively. *Tg*(*X1a.Eef11:H2B-mCherry*) line to visualise nuclei was previously described [26]. *Tg*(*Ubi::RacFRET*) line was generated in this study.

### DNA constructs, mRNA synthesis, morpholinos and injection procedures

The transgenic p*Tol2-Ubiquitin-RacFRET-βGlobin* construct for was created by subcloning the *RacFRET-NoCT* sequence [14]) into pTol2 plasmid containing ubiquitin promoter kindly provided by Sebastian Gerety [27]. A linearised DNA encoding for EGFP-Rac1-Globin was used as a template for mRNA synthesis to produce EGFP-Rac1 in rescue experiments. *EGPF-Rac1* was used as a control mRNA in rescue experiments.

Capped sense RNA was *in vitro* transcribed using the mMessage mMachine kit (Thermo Fisher Scientific). The mRNA was purified using RNeasy columns from Qiagen. The purified sense mRNA was injected into one-cell stage embryos, 300pg per embryo. To inhibit protein translation, morpholino antisense oligonucleotides (Gene Tools) were injected into one-cell stage embryos. *rac1a* splice morpholino: *5’CCACACACTTTATGGCCTGCATCTG3’* [15]; *rac2* morpholino: *5’CCACCACACACTTTATTGCTTGCAT3’* [17]; *Standard morpholino: 5’CCTCTTACCTCAGTTACAATTTATA3’*. Morphonlinos were injected at 1.5 pmol per embryo.

### Phalloidin staining

The whole mount phalloidin staining was done according to the protocol [28]. In brief, the embryos were fixed with 4% PFA in PBS overnight at 4ºC. The next day, the embryos were washed with PBS-Tween (0.1%) three times of 5-minute intervals and incubated with phalloidin 488 diluted 1:20 in PBS-Tween (0.1%) overnight at 4ºC. Prior imaging, the embryos were washed with PBS-Tween (0.1%) and kept in PBS during imaging.

### Imaging pectoral fin growth

Fin growth was monitored on the upright Zeiss LSM 710 equipped with 20x water immersion objective (NA 1) using one-photon mode. The embryos were kept at 28°C and the chorion was removed at 24 hpf using forceps. To image fin growth, the embryos were anaesthetised with 0.04% of tricaine methanesulfonate (Sigma) in embryo medium and mounted dorsally using agarose moulds. The embryo was pushed into the rectangular slot made in the agarose (approximately 0.5 mm width and depth) in such manner that the yolk was slightly compressed by the agarose walls while exposing the embryo with its dorsal side facing the objective. Such mounting immobilised the embryo while allowing pectoral fin to develop unperturbed. The embryo was tilted slightly to the right or to the left to allow the confocal scanning plane parallel to the plane formed by AP and DV axes. To validate the results, each experiment was repeated at least three times on different days.

### Laser ablation

Laser ablation was performed on spinning disk setup equipped with UV laser as described in [24, 29]. Briefly, the double transgenic embryos expressing Lifeact-GFP and Lyn-Tomato under the control of the *actb1* promoter, were mounted on a spinning-disk microscope equipped with 355nm UV laser. 63W x 1.2 NA objective was used. The ablation was done by pulsing the laser along single line perpendicular to the orientation of the actin belt (AP), pulse per shot: 50, shots per micron: 2.0. The delay time: 0.725 s.

### Image processing

Image analysis and quantifications were done using Fiji/ImageJ software.

#### Measuring fin dimensions

Fin dimensions were obtained by measuring the length along the AP, DV and PD axes.

When necessary, the xyz drift was corrected by the image registration plugin: [Plugins > Registration > Correct 3D drift].

To obtain the AP and DV dimensions, the following steps were followed:

- [Image > Stacks > Z Project …] (‘Sum Slices’ or ‘Standard Deviation’ modes).
- Segment the image: [Image > Adjust > Threshold].
- Select the area using ‘magic wand’ tool.
- [Edit > Selection > Fit Ellipse].
- The major and minor axes of the ellipse correspond to the AP and DV fin axes respectively.
- Obtain AP and DV dimensions after applying: [Image > Stacks > Reslice[/] …] command along the AP or DV axes respectively.

To obtain cross-section along AP or DV axis for the lateral and sagittal views respectively, the Z-stack was dissected along the AP or DV axes using: [Image>Stacks>Reslice[/]…] command.

To measure the PD axis dimensions, the fin was optically dissected along the AP axis using: [Image > Stacks > Reslice[/].] command and the maximal length was obtained by measuring the distance between the proximal and distant.

To measure the actin belt length: the optical section along the AP axis was used to obtain the arch or acting belt and the length was measured manually by fitting a segmented line into the actin belt.

#### Cell tracking

Cells were tracked using Manual Tracking plugin of Fiji: [Plugins > Tracking > Manual Tracking].

#### Cell number counting

To obtain cell number in the fin at 48 hours, cells were counted manually by labelling each nucleus in the image using the ‘brush’ tool in Fiji. After every cell was labelled, the mask was applied to convert image to a binary format: [Process > Bianry > Make Binary] followed by the [Analyze > Analyze Particles.] command was used to obtain the number of cells. To quantify fin dimensions and cell number, at least 5 embryos were used.

Rac activity was analysed by measuring FRET ratio as previously described [14].

### Statistical analysis

Statistical analyses and graphs were generated with Excel or GraphPad Prism software. *p* values indicated on Fig. 3B and C correspond to Student’s t tests, which have been performed after the normality of the distributions studied was verified by generating normal probability plots (P-P plots).

## Supporting information

Supplemental movies list

## Acknowledgements and funding

This work was supported by the long-term FEBS postdoctoral fellowship to E.K. between 2012 and 2014; epiPhysX S18011 between 2014 and 2016; and ANR-10-INBS-04 through the National Infrastructure France-BioImaging supported by the French National Research Agency and ITN H2020 ImageInLife between 2017-2018. H.N is supported by 2017-ITN-721537 as part of the ITN ImageInLife Marie Skłodowska-Curie Actions. The authors thank Rozenn Finazzi and Susanne Borgers for technical assistance.

---

**Supplemental Figure 1.**
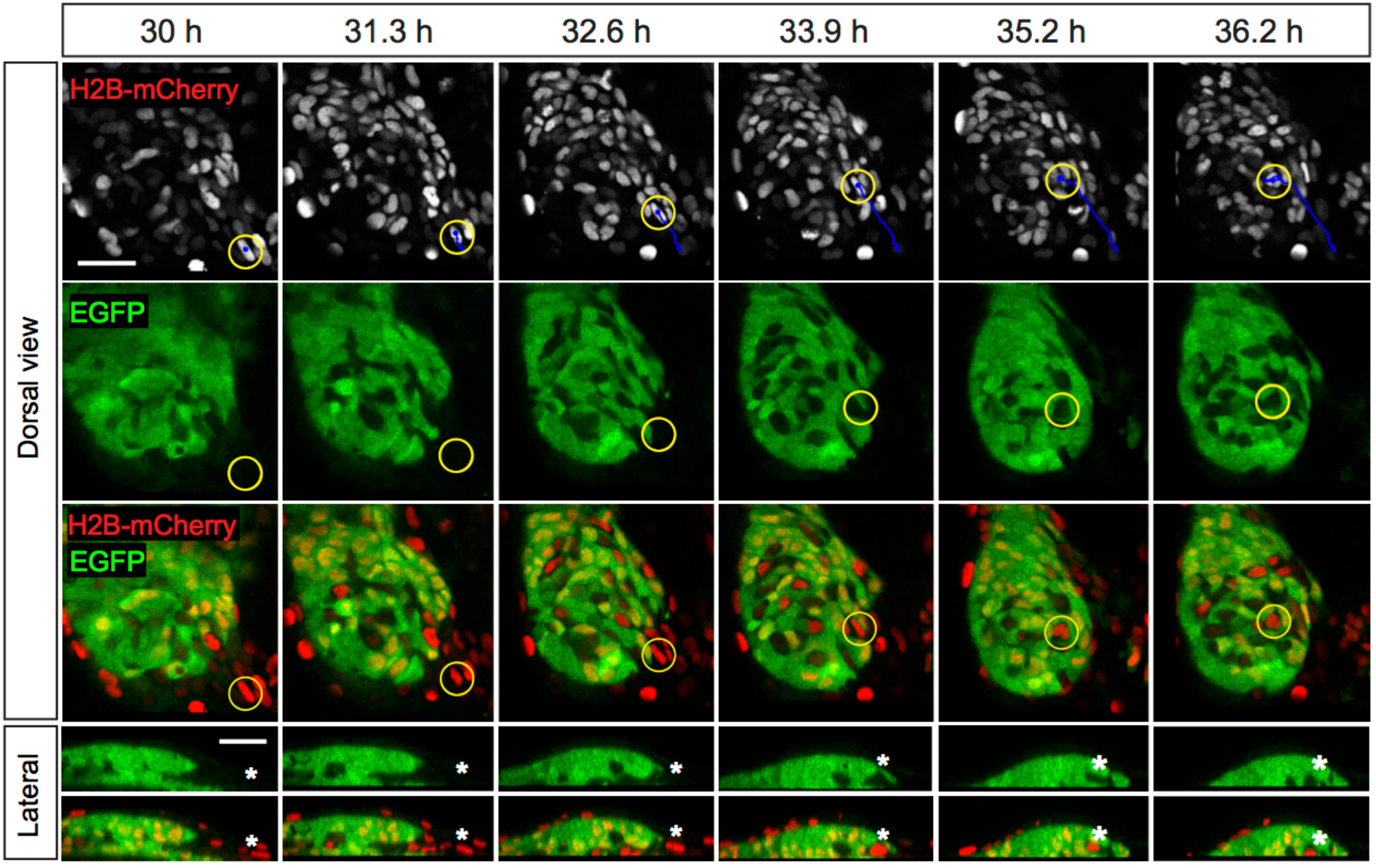
Cell migration into the fin field from the area neighboring the 4^th^ somite. *Tg*(*mPrx1(cFos):EGFP*) embryos were injected at one-cell stage with mRNA encoding for H2B-mCherry. A single confocal section with 0.8μm thickness for each time frame is shown for the dorsal view. The EGFP-negative cell migrating into the fin area is marked with a yellow circle and the tracking path is shown in blue. The lateral panel corresponds to the cross-section along the cell migratory path and the apteryx marks the location of the invading cell. Scale bar: 30μm. See also Supplemental movie 3.

**Supplemental Figure 2.**
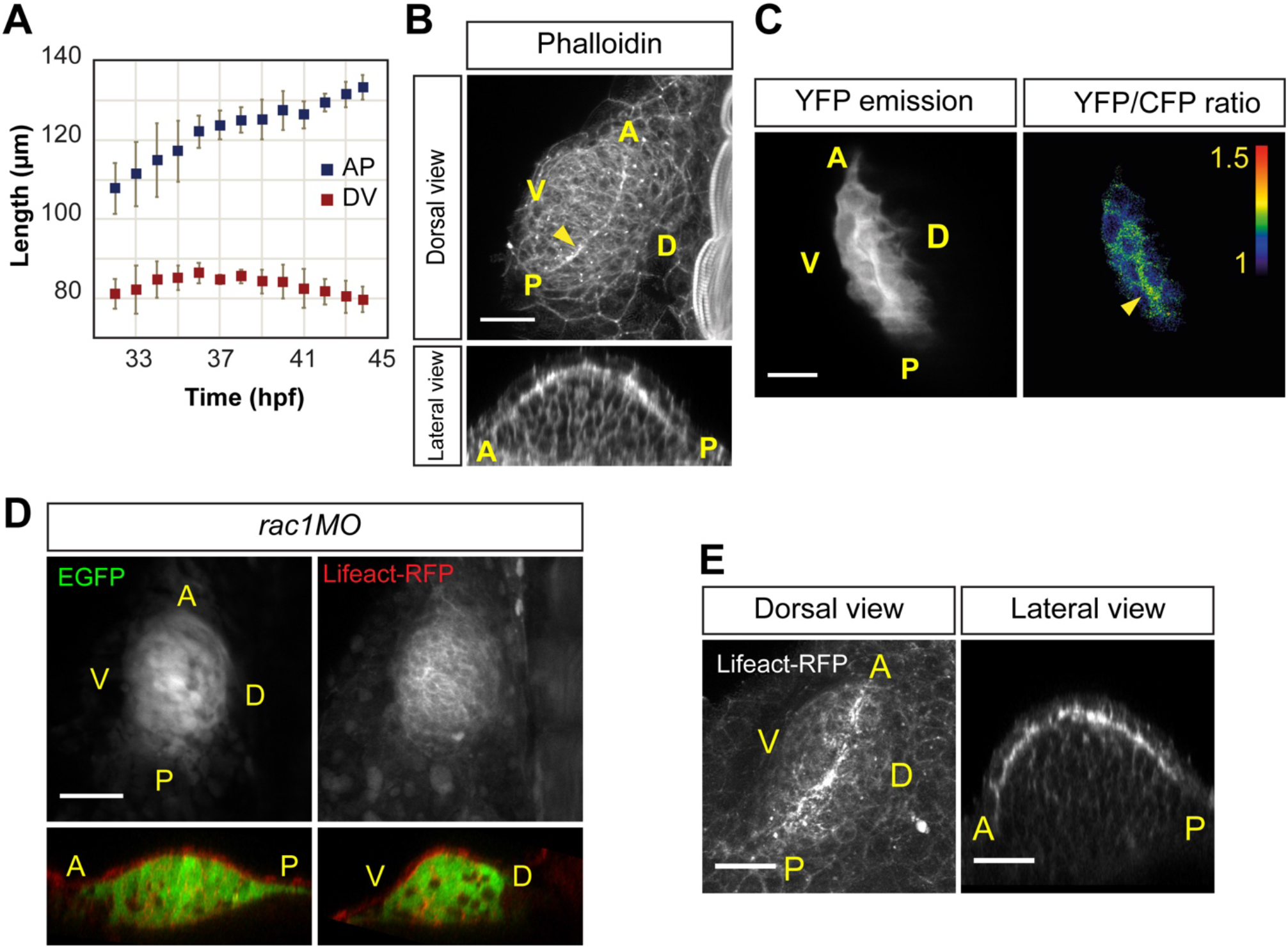
Actin belt dynamics and fin shape regulation. **A.** The graph shows the dynamics of AP and DV fin dimensions in *Tg*(*mPrx1(cFos):EGFP*) embryos between 32 and 44 hpf. N = 6. Data are represented as mean ± standard deviation. See also Figure 1D. **B.** Phalloidin staining of the endogenous actin in the pectoral fin at 48 hpf. Wild type embryos were used and the representative fin is shown. Scale bar: 30μm. **C.** Rac activity in the ectodermal layer of the pectoral fin at the dorso-ventral boundary is measured in a *Tg*(*Ubi::RacFRET*) embryo at 46 hpf. Scale bar: 20μm. **D.** Pectoral fin in a double transgenic *Tg*(*mPrx1(cFos):EGFP); Tg(actb1:lifeact-RFP*) embryos treated with *rac1* morpholino at 46 hpf. Scale bar: 40μm. **E.** Rescue of the actin belt formation by the over-expression of EGFP-Rac 1 in *Rac1* morpholino-treated transgenic *Tg*(*actb1:lifeact-RFP*) embryos. A representative pectoral fin in the embryo at 46 hpf. Scale bar: 30μm.

**Supplemental Figure 3.**
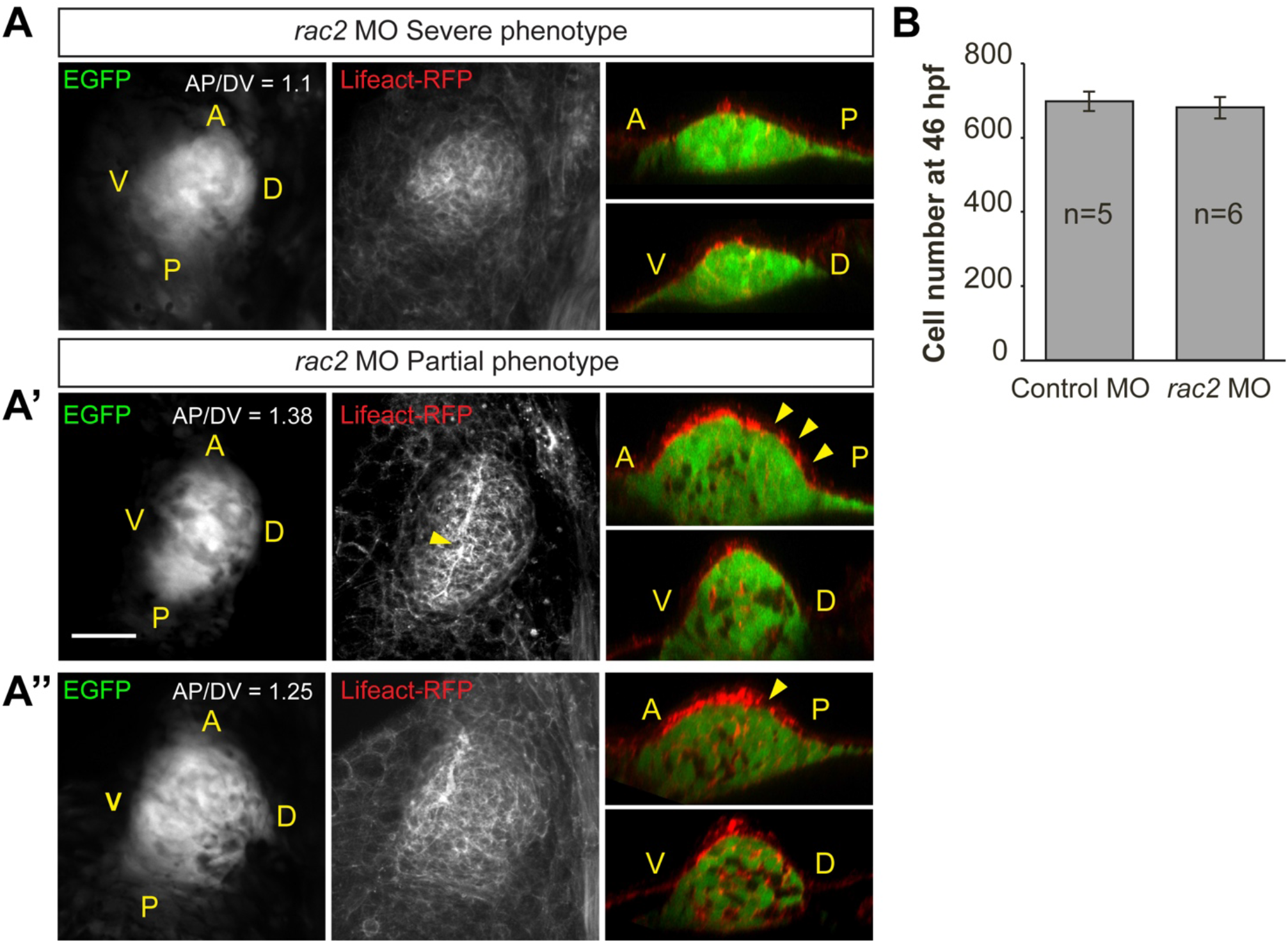
Rac2 morphant phenotype categories classification. Examples for partial and severe phenotypes in pectoral fins of double transgenic embryos *Tg*(*mPrx1(cFos):EGFP*); *Tg*(*actb1:Lifeact-RFP*) treated with *rac2* morpholinos. **A.** A representative fin with the severe phenotype: the actin belt is missing and the fin is smaller in size. **A’ and A’’** show two typical fins exhibiting the partial phenotype: the actin belt is partially formed and the shape is affected. Scale bar: 40μm. AP/DV ratio is indicated for each fin. **B.** The graph compares cell number in the pectoral fins at 46 hpf between the control and *rac2* morpholino treated embryos. Only the embryos belonging to the “partial phenotype” category were selected for this analysis. Data are represented as mean ± SEM. See also Figure 3.

**Supplemental Figure 4.**
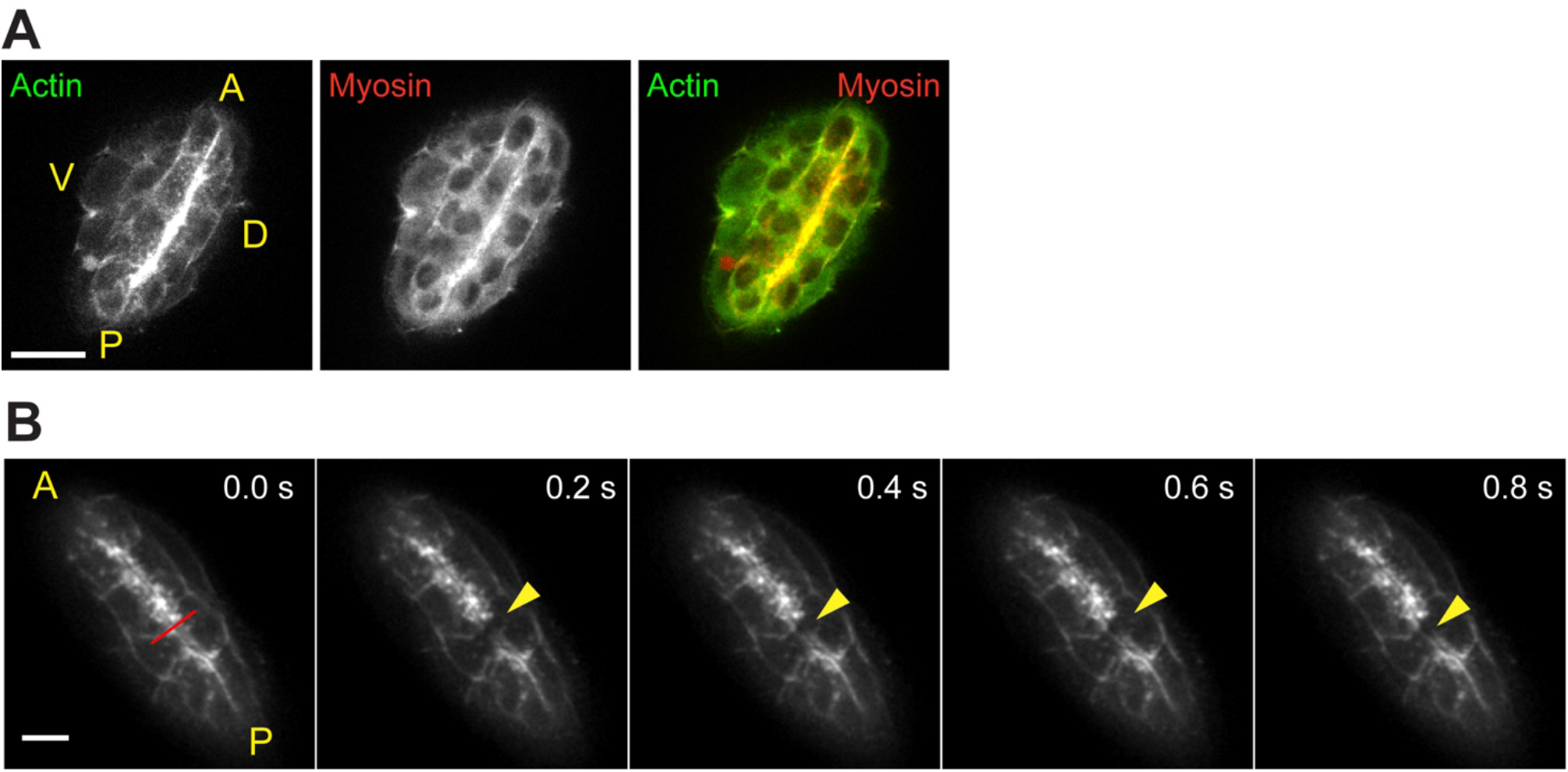
Actin belt contractility in the pectoral fin at 46 hpf. **A.** Confocal sections of the distal portion of the fin in a double transgenic embryo *Tg*(*actb1-lifeact-GFP*); *Tg*(*β-actin::myl12.1-mCherry*). Scale bar: 20μm. **B.** The actin belt in the pectoral fin of the transgenic embryo *Tg*(*actb1:lifeact-GFP*) is under tension at 48. The red line indicates the direction of the laser cut and is perpendicular to the AP axis of the fin and the orientation of the actin belt. A yellow arrow points at the recoil induced by the laser cut. Scale bar: 10 μm. See also the Supplemental Movie 10.

## References

1. Mercader N. (2007). Early steps of paired fin development in zebrafish compared with tetrapod limb development. Development, Growth & Differentiation 49, 421–437.

2. Ahn, D.-g., Kourakis, M.J., Rohde, L.A., Silver, L.M., and Ho, R.K. (2002). T-box gene tbx5 is essential for formation of the pectoral limb bud. Nature 417, 754–758.

3. Yano, T., Abe, G., Yokoyama, H., Kawakami, K., and Tamura, K. (2012). Mechanism of pectoral fin outgrowth in zebrafish development. Development (Cambridge, England) 139, 2916–2925.

4. Masselink, W., Cole, N.J., Fenyes, F., Berger, S., Sonntag, C., Wood, A., Nguyen, P.D., Cohen, N., Knopf, F., Weidinger, G., et al. (2016). A somitic contribution to the apical ectodermal ridge is essential for fin formation. Nature, 1–16.

5. Mao, Q., Stinnett, H.K., and Ho, R.K. (2015). Asymmetric cell convergence-driven zebrafish fin bud initiation and pre-pattern requires Tbx5a control of a mesenchymal Fgf signal. Development (Cambridge, England) 142, 4329–4339.

6. Charité, J., McFadden, D.G., and Olson, E.N. (2000). The bHLH transcription factor dHAND controls Sonic hedgehog expression and establishment of the zone of polarizing activity during limb development. In Development, Volume 127. pp. 2461–2470.

7. Galli, A., Robay, D., Osterwalder, M., Bao, X., Bénazet, J.-D., Tariq, M., Paro, R., Mackem, S., and Zeller, R. (2010). Distinct roles of Hand2 in initiating polarity and posterior Shh expression during the onset of mouse limb bud development. In PLoS Genet, Volume 6. p. e1000901.

8. te Welscher, P., Fernandez-Teran, M., Ros, M.A., and Zeller, R. (2002). Mutual genetic antagonism involving GLI3 and dHAND prepatterns the vertebrate limb bud mesenchyme prior to SHH signaling. In Genes & Development, Volume 16. pp. 421–426.

9. Hernández-Vega, A., and Minguillón, C. (2011). The Prx1 limb enhancers: Targeted gene expression in developing zebrafish pectoral fins. Developmental dynamics : an official publication of the American Association of Anatomists 240, 1977–1988.

10. Zeller, R., López-Ríos, J., and Zuniga, A. (2009). Vertebrate limb bud development: moving towards integrative analysis of organogenesis. Nature Reviews Genetics 10, 845–858.

11. Nacu, E., and Tanaka, E.M. (2011). Limb regeneration: a new development? Annual Review of Cell and Developmental Biology 27, 409–440.

12. Lau, K., Tao, H., Liu, H., Wen, J., Sturgeon, K., Sorfazlian, N., Lazic, S., Burrows, J.T.A., Wong, M.D., Li, D., et al. (2015). Anisotropic stress orients remodelling of mammalian limb bud ectoderm. Nature Cell Biology 17, 569–579.

13. Haga, R.B., and Ridley, A.J. (2016). Rho GTPases: Regulation and roles in cancer cell biology. Small GTPases 7, 207–221.

14. Kardash, E., Bandemer, J., and Raz, E. (2011). Imaging protein activity in live embryos using fluorescence resonance energy transfer biosensors. Nature Protocols 6, 1835–1846.

15. Srinivas, B.P., Woo, J., Leong, W.Y., and Roy, S. (2007). A conserved molecular pathway mediates myoblast fusion in insects and vertebrates. In Nat Genet, Volume 39. pp. 781–786.

16. Thisse, B., Thisse, C. (2004). Fast Release Clones: A High Throughput Expression Analysis..

17. Deng, Q., Yoo, S.K., Cavnar, P.J., Green, J.M., and Huttenlocher, A. (2011). Dual roles for Rac2 in neutrophil motility and active retention in zebrafish hematopoietic tissue. In Developmental Cell, Volume 21. pp. 735–745.

18. Rosowski, E.E., Deng, Q., Keller, N.P., and Huttenlocher, A. (2016). Rac2 Functions in Both Neutrophils and Macrophages To Mediate Motility and Host Defense in Larval Zebrafish. In The Journal of Immunology, Volume 197. pp. 4780–4790.

19. Wyngaarden, L.A., Vogeli, K.M., Ciruna, B.G., Wells, M., Hadjantonakis, A.-K., and Hopyan, S. (2010). Oriented cell motility and division underlie early limb bud morphogenesis. Development (Cambridge, England) 137, 2551–2558.

20. Hopyan, S., Sharpe, J., and Yang, Y. (2011). Budding behaviors: Growth of the limb as a model of morphogenesis. Developmental dynamics : an official publication of the American Association of Anatomists 240, 1054–1062.

21. Marcon, L., Arqués, C.G., Torres, M.S., and Sharpe, J. (2011). A Computational Clonal Analysis of the Developing Mouse Limb Bud. PLoS Computational Biology 7, e1001071.

22. Legoff, L., Rouault, H., and Lecuit, T. (2013). A global pattern of mechanical stress polarizes cell divisions and cell shape in the growing Drosophila wing disc. Development (Cambridge, England) 140, 4051–4059.

23. Krens, S., and Heisenberg, C.-P. (2011). Cell sorting in development. Current topics in developmental biology 95, 189–213.

24. Behrndt, M., Salbreux, G., Campinho, P., Hauschild, R., Oswald, F., Roensch, J., Grill, S.W., and Heisenberg, C.-P. (2012). Forces driving epithelial spreading in zebrafish gastrulation. Science 338, 257–260.

25. Maître, J.-L., Berthoumieux, H., Krens, S.F.G., Salbreux, G., Jülicher, F., Paluch, E., and Heisenberg, C.-P. (2012). Adhesion functions in cell sorting by mechanically coupling the cortices of adhering cells. Science 338, 253–256.

26. Recher, G., Jouralet, J., Brombin, A., Heuzé, A., Mugniery, E., Hermel, J.-M., Desnoulez, S., Savy, T., Herbomel, P., Bourrat, F., et al. (2013). Zebrafish midbrain slow-amplifying progenitors exhibit high levels of transcripts for nucleotide and ribosome biogenesis. Development (Cambridge, England) 140, 4860–4869.

27. Gerety, S.S., Breau, M.A., Sasai, N., Xu, Q., Briscoe, J., and Wilkinson, D.G. (2013). An inducible transgene expression system for zebrafish and chick. Development (Cambridge, England) 140, 2235–2243.

28. Goody, M.F., and Henry, C.A. (2013). Phalloidin Staining and Immunohistochemistry of Zebrafish Embryos. Bio-protocol 3, e786.

29. Smutny, M., Behrndt, M., Campinho, P., Ruprecht, V., and Heisenberg, C.-P. (2015). UV laser ablation to measure cell and tissue-generated forces in the zebrafish embryo in vivo and ex vivo. Methods in molecular biology (Clifton, NJ) 1189, 219–235.

