## Supplemental movies list for "A contractile acto-myosin belt promotes growth anisotropy during the early stages of pectoral fin development in zebrafish"

### Supplemental Information

#### Supplemental Movies list and access links:

**Supplemental Movie 1.** Cellular dynamics between 32 and 35 hpf in the transgenic embryo *Tg(X1a.Eef1α:H2B-mCherry*) reveals cells invading fin field from the region neighbouringåœåå 4th somite.
https://www.dropbox.com/s/2ogyprmgrmyb7d0/S1_migration_32h.mov?dl=0
**Supplemental Movie 2.** Cellular dynamics between 34 and 42 hpf in the transgenic embryo *Tg(X1a.Eef1α:H2B-mCherry*) reveals cells invading the fin field from the region neighbouring 4th somite.
https://www.dropbox.com/s/72luhu3s1ckyorz/S2_migration_34h.mov?dl=0
**Supplemental Movie 3.** The EGFP-negative cell migrates into the fin area. This movie corresponds to supplemental Figure 1. *mPrx1(cFos):EGFP* embryos were injected at one-cell stage with mRNA encoding for H2B-mCherry. A Z projection of the confocal sections from z-tack covering the path of a migrating cells were used to generate this movie. The total thickness of a section is 8.8 $\mu$m. Scale bar: 30$\mu$m.
https://www.dropbox.com/s/gjrcrgxk0en1kec/S3_Migrate_Tracked.mov?dl=0
**Supplemental Movie 4.** 3D fin shape at 28 hpf in a *Tg*(*mPrx1(cFos):EGFP)* embryo.
https://www.dropbox.com/s/5rhv48gntbsxkkv/S4_3D_28h.mov?dl=0
**Supplemental Movie 5.** 3D fin shape at 46 hpf in a *Tg*(*mPrx1(cFos):EGFP)* embryo.
https://www.dropbox.com/s/waxprglnf5o9ei0/S5_3D_46h.mov?dl=0
**Supplemental Movie 6.** A confocal Z projection along the proximal-distal axis of the fin at 46 hpf in a transgenic embryo expressing Lifeact-GFP in the fin.
https://www.dropbox.com/s/8xqy7wtggrcjods/S6_Actin02lmaG2_130613.mov?dl=0
**Supplemental Movie 7.** Actin belt formation in a transgenic embryo expressing Lifeact-EGFP in the fin.
https://www.dropbox.com/s/r9jmir5ia46jbdq/S7_Actin_Belt.avi?dl=0
**Supplemental Movie 8.** A 3D reconstruction of the fin at 50 hpf in the double transgenic embryo expressing H2B-mCherry and Lifeact-EGFP in the fin.
https://www.dropbox.com/s/pu4e0z084vh54g8/S8_Actin3D-56h.mov?dl=0
**Supplemental Movie 9.** Cellular dynamics in the *rac2* morphant fins. The *Tg(X1a.Eef1α:H2B-mCherry*) embryos were injected with *rac2* morpholino at one cell stage.
https://www.dropbox.com/s/1q56924r187x2rv/S9_Rac2MO.mov?dl=0
**Supplemental Movie 10.** Laser ablation of the actin belt in the fin at 46 hpf in the *Tg(actb1:lifeact-GFP*) embryo. https://www.dropbox.com/s/rvlxwp0i1968t0b/S10_Ablation.mov?dl=0
